# Multi-omic profiling of the leukemic microenvironment shows bone marrow interstitial fluid is distinct from peripheral blood plasma

**DOI:** 10.1101/2022.03.30.486272

**Authors:** Lorenz Nierves, Jian Guo, Siyuan Chen, Janice Tsui, Anuli Uzozie, Jonathan W. Bush, Tao Huan, Philipp F. Lange

**Author notes:** Correspondence to: Philipp F. Lange.

## Abstract

The bone marrow is the place of hematopoiesis with a microenvironment that supports lifelong maintenance of stem cells and high proliferation. It is not surprising that this environment is also favourable for malignant cells emerging in the bone marrow or metastasizing to it. While the cellular composition of the bone marrow microenvironment has been extensively studied, the extracellular matrix and interstitial fluid components have received little attention. Since the sinusoids connect the bone marrow interstitial fluid to the circulation, it is often considered to have the same composition as peripheral blood plasma. Stark differences in the cellular composition of the bone marrow and peripheral blood with different secretory capacities would however suggest profound differences.

In this study we set out to better define if and how the bone marrow interstitial fluid (BMIF) compares to the peripheral blood plasma (PBP) and how both are remodeled during chemotherapy. We applied a multi-omic strategy to quantify the metabolite, lipid and protein components as well as the proteolytic modification of proteins to gain a comprehensive understanding of the two compartments. We found that the bone marrow interstitial fluid is clearly distinct from peripheral blood plasma, both during active pediatric acute lymphoblastic leukemia and following induction chemotherapy. Either compartment was shaped differently by active leukemia, with the bone marrow interstitial fluid being rich in extracellular vesicle components and showing protease dysregulation while the peripheral blood plasma showed elevation of immune regulatory proteins. Following chemotherapy, the BMIF showed signs of cellular remodeling and impaired innate immune activation while the peripheral blood plasma was characterized by restored lipid homeostasis.

## 1. INTRODUCTION

The tumour microenvironment — comprised of stromal cells, extracellular matrix, and interstitial fluid — plays a central role in the maintenance of cancer. In blood cancers, including pediatric acute lymphoblastic leukemia (ALL), the bone marrow and the peripheral blood are the two primary microenvironments and much progress has been made in understanding them. Leukemic cell growth is dependent on the microenvironment [1] and, in turn, cancer cells outcompete normal cells, thereby disrupting and remodeling the microenvironment to further drive malignant cell expansion [2,3]. While we now have detailed understanding of the role of individual cellular components, such as stromal or immune cell populations [4,5], the fluid portion of the microenvironment is often overlooked and studies that explore it have been scant.

The complexity and wide range of analyte abundancies spanning several orders of magnitude pose challenges to the comprehensive characterization of both compartments. But while the proteomic investigation of blood plasma has seen major advances [6,7], a comprehensive investigation into these ALL-relevant biofluids is lacking. Previously, the leukemic BMIF was profiled using a cytokine antibody array to compare BMIF collected at diagnosis and after bone marrow transplant to identify tumourigenic soluble proteins [8]. In a separate study, the leukemic and normal PBP was profiled by proteomics [9].

Although there is a path for migration between the bone marrow and the peripheral blood, the extent of such movement on the BMIF and the PBP has not been fully explored. Initial evidence from previous studies, however, provide basis for our hypothesis that the BMIF and the PBP are different. BMIF collected from patients with hematologic disorders had lower levels of select metabolites and proteins compared to corresponding plasma [10]. Furthermore, *in vitro* characterization of BMIF and PBP from adult acute myeloid leukemia (AML) patients showed differences in the potential to inhibit hematopoietic progenitor cell growth linked to different levels of TNF*α* and adiponectin [11]. The study further showed that induction chemotherapy leads to a drop in TNF*α* and adiponectin in BMIF while their concentration remained unchanged in the peripheral blood. This, as well as documented differences in cellular composition [12], suggests functionally important differences in the composition of peripheral blood plasma and bone marrow interstitial fluid.

To investigate if bone marrow interstitial fluid and peripheral blood plasma differ by more than a few select chemokines, we employed a multi-omic approach to characterize both fluid portions of the ALL microenvironment. We analyzed the proteome, protein processing, metabolome and lipidome in a retrospective collection of BMIF and PBP from 8 pediatric B-ALL patients. This first comprehensive comparison of BMIF and PBP demonstrated widespread differences between the two compartments that span molecular classes and manifest in a compartment specific response to therapy.

## 2. RESULTS AND DISCUSSION

### 2.1. Bone marrow interstitial fluid and peripheral blood plasma have distinct proteome and metabolome landscapes

To investigate the molecular characteristics and relationships of the two ALL-relevant microenvironments, and to understand their remodeling during chemotherapy, we collected bone marrow interstitial fluid (BMIF) and peripheral blood plasma (PBP) from 8 pediatric B-cell acute lymphoblastic leukemia (B-ALL) patients (Supplementary Table 1). As expected, peripheral blood and bone marrow smears show that the peripheral blood and bone marrow compartments differ in their cell composition (Supplementary Figure 1). The bone marrow also shows the expected changes in cellular composition following blast eradication (Figure 1A). To investigate if these structural and cellular differences lead to differences in the molecular composition, matched BMIF and PBP collected from the same patient at time of diagnosis and after the induction phase of chemotherapy (Figure 1B) were subjected to four sample processing and mass spectrometry-based strategies to obtain quantitative proteomic, terminomic, metabolomic and lipidomic profiles (Figure 1C).

**Figure 1.**
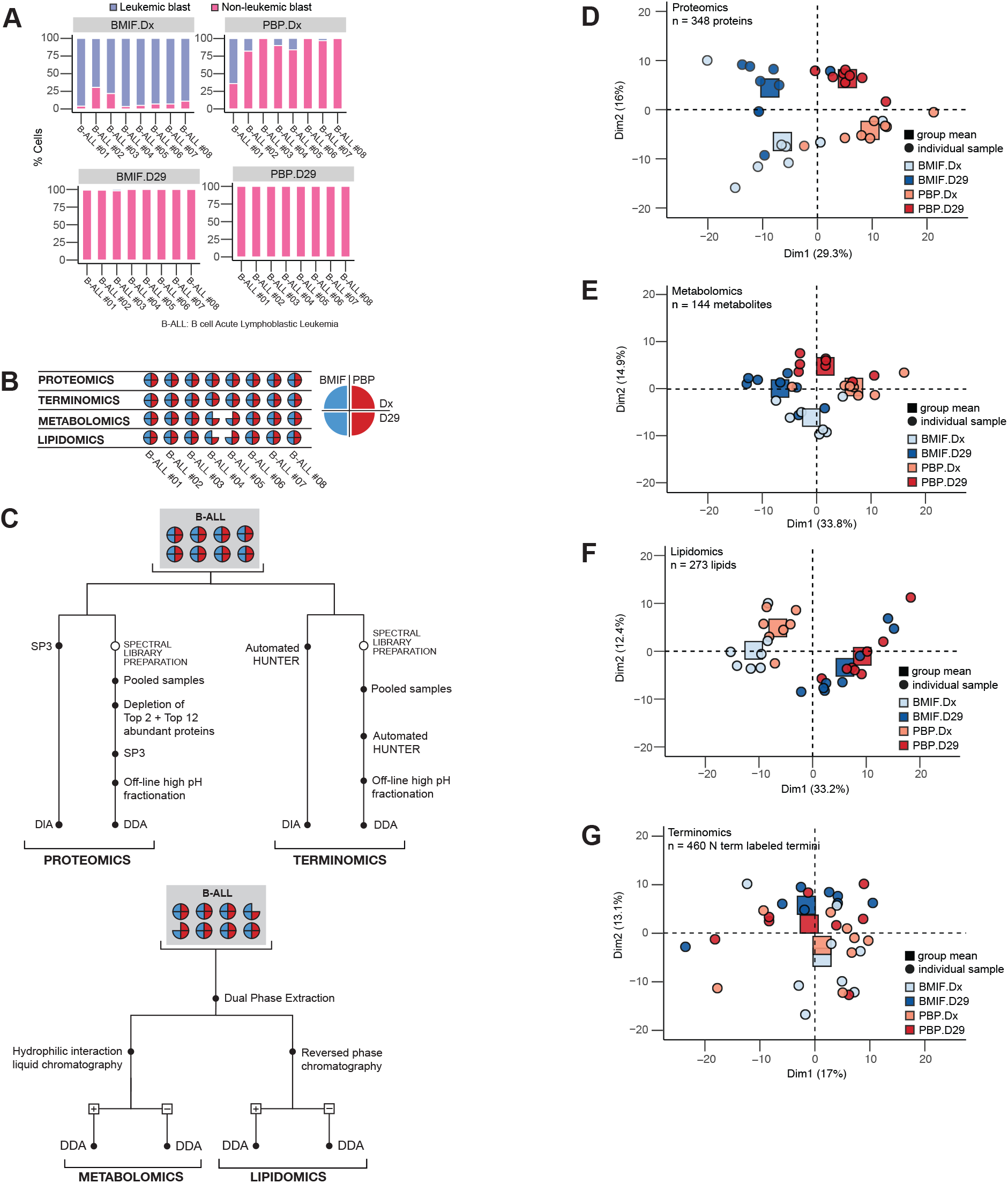
The BMIF and the PBP are molecularly distinct. **(A)** Cellular composition of the BMIF and PBP compartments at Dx and D29. Stacked bars represent the proportion of leukemic and non-leukemic cell blasts in each B-ALL patient. **(B)** Schematic summarizing specimen collection and analyses. Bone marrow interstitial fluid (BMIF) and peripheral blood plasma (PBP) were collected at diagnosis (Dx) and after post-induction therapy (D29) for 8 pediatric B-ALL patients. Samples were processed for proteomic, terminomic, metabolomic, and lipidomic analyses. B-ALL #04 and B-ALL #05, did not have sufficient material for metabolomic and lipidomic analysis. **(C)** Sample preparation workflows for multi-omic mass spectrometric analysis. (D-G) Principal Component Analysis (PCA) of **(D)** proteomics **(E)** metabolomics **(F)** lipidomics, and **(G)** terminomics data. Individual samples are plotted as circles and the mean principal components of experimental group means are plotted as squares.

Proteomic and terminomic analyses were conducted using Single-pot solid-phase-enhanced sample preparation (SP3) [13] and High-efficiency Undecanal-based N Termini EnRichment (HUNTER) [14], respectively. Extracted analytes were analysed by LC-MS/MS using Data Independent Acquisition (DIA). We quantified a total of 528 protein groups across all patients, with 415 (79%) quantified in all experimental groups (Supplementary Figure 2A, Supplementary Figure 2B). The quantified proteins spanned 6 orders of magnitude (Supplementary Figure 2C) and were primarily associated with extracellular annotation based on COMPARTMENTS [15] (Supplementary Figure 2D).

**Figure 2.**
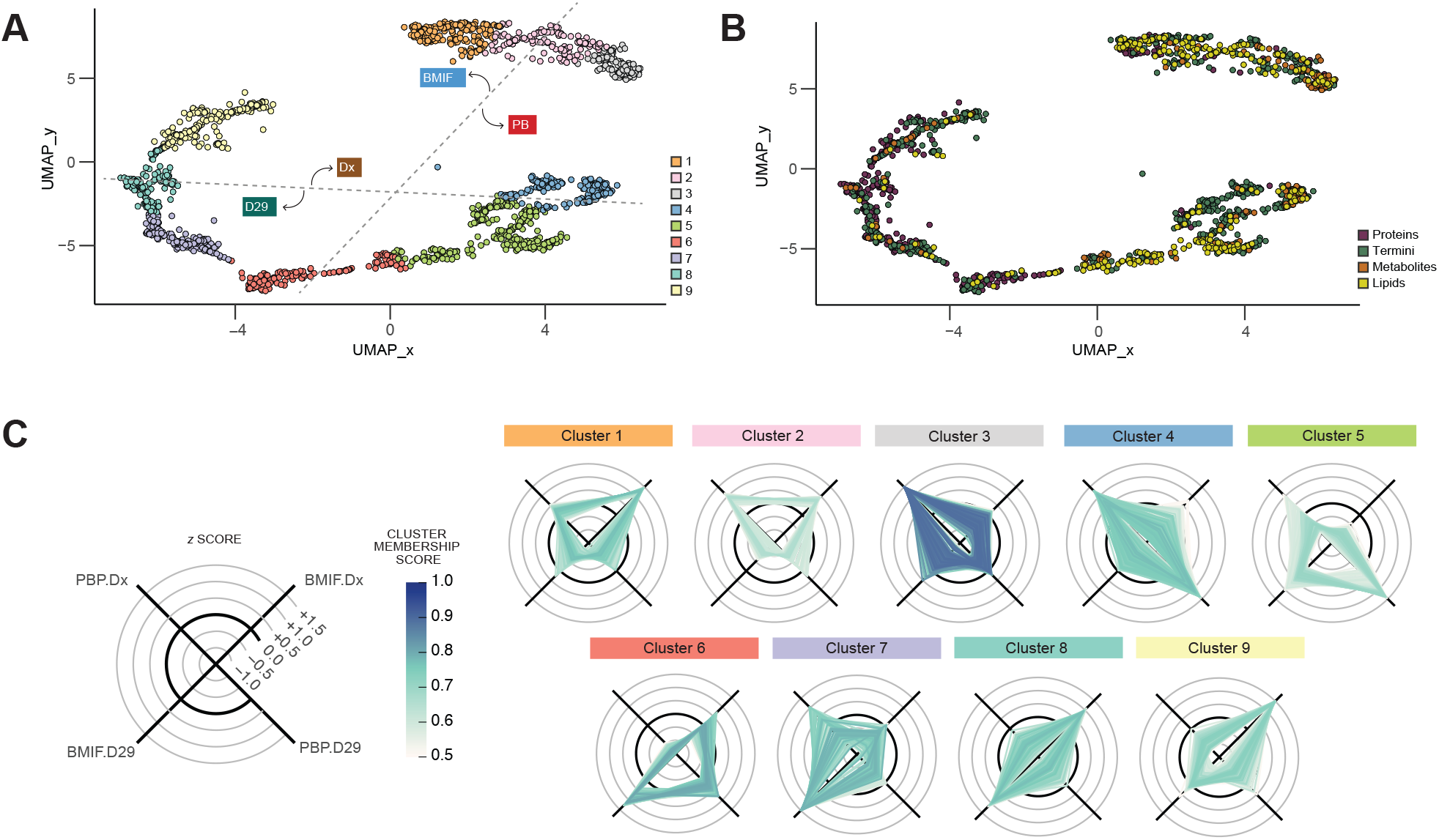
Integration of data reveals that multiple analytes differ between BMIF and PBP. Uniform Manifold Approximation and Projection (UMAP) of all analytes (normalized and z-score scaled) after training using significant analytes. Individual analytes are annotated based on **(A)** fuzzy c-means cluster membership or **(B)** analyte class. For the cluster membership plot, dotted lines were draw to indicate separation between experimental groups. **(C)** Radar plots showing intensity of the analytes in each cluster. Z score-scaled intensities were plotted and annotated based on the analyte’s cluster membership score. Color intensity reflects the cluster score and analytes with scores <0.5 are omitted.

Using HUNTER we identified 915 unique N termini, representing 201 proteins (Supplementary Figure 2E) spanning 7 orders of magnitude (Supplementary Figure 2F). As with SP3 based protein identifications, proteins that were identified by their N termini using the HUNTER workflow, were also primarily located in the extracellular space (Supplementary Figure 2G). We estimated the concentration range captured in our study based on concentrations (mostly from adults) reported by the Human Protein Atlas database (proteinatlas.org) [16]. Of the 588 proteins detected by SP3 and HUNTER, we were able to match the annotated plasma concentration for 403 proteins. The detected proteins in our data were from a wide concentration range spanning 8 orders of magnitude from 830mg/L (CP) to 8.9ng/L (PLIN1) (Supplementary Figure 1H).

Metabolomics and lipidomics data were obtained by processing the patient samples via Dual Phase Extraction [17] followed by HILIC and Reversed Phase chromatography and MS acquisition in positive and negative mode (Figure 1B). In total, we quantified 160 unique metabolites (Supplementary Figure 3A) and 279 unique lipids across 5 lipid classes (Supplementary Figure 3B) spanning 5 and 4 orders of magnitude respectively (Supplementary Figures 3C, D). 150 (94%) metabolites and all 279 lipids were represented in all four experimental groups (Supplementary Figures 3E, F).

**Figure 3.**
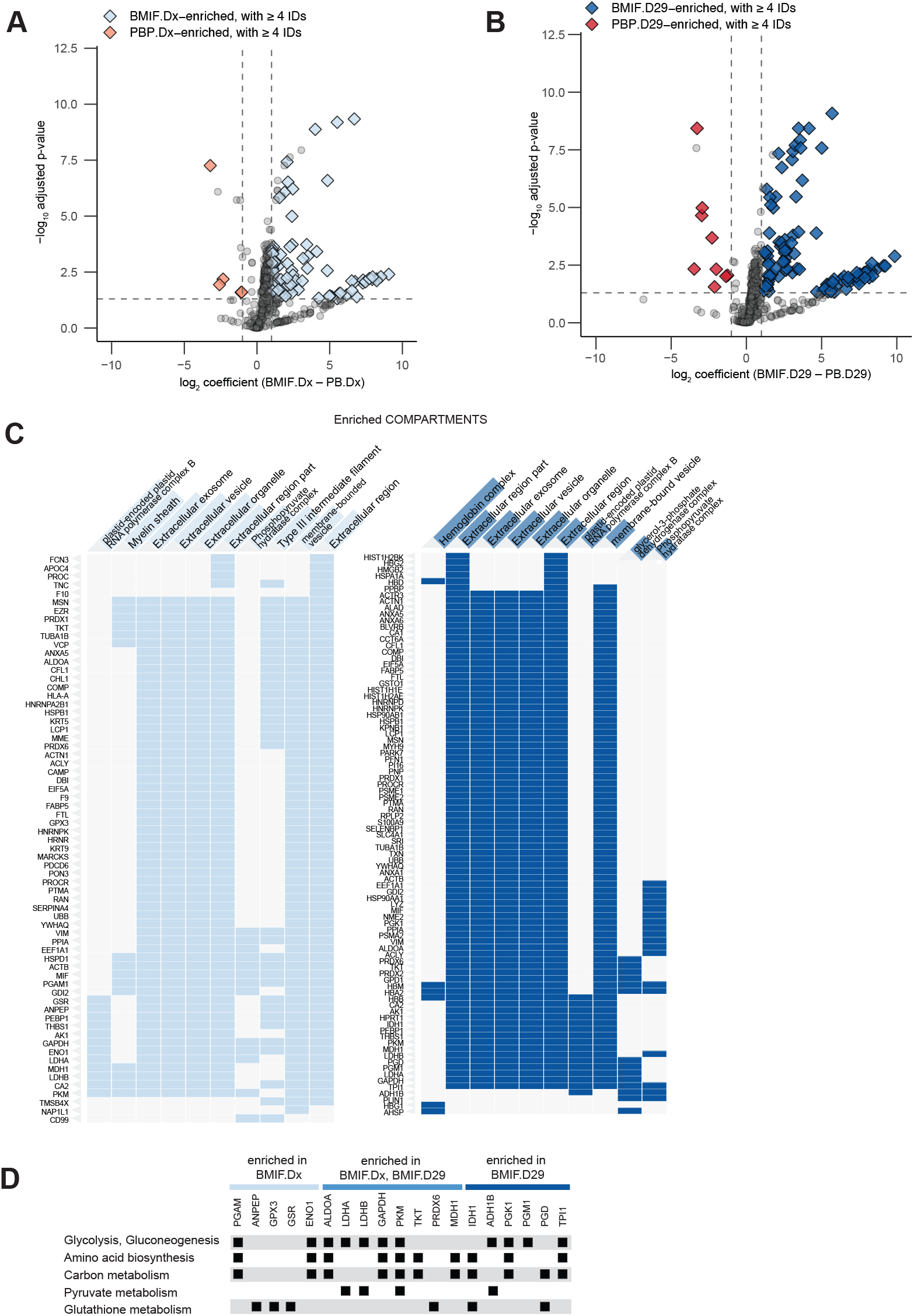
Proteins enriched in the BMIF are localized in extracellular vesicles and exosomes. **(A)** Compartment differences at leukemia (BMIFdx vs PBPdx). **(B)** Compartment differences after induction therapy (BMIF.D29 vs PBP.D29). Both volcano plots show –log10 adjusted p-value and log2 coefficient of each protein after limma analysis. Dashed lines mark the cut-offs for significance in their respective intercepts. **(C)** Subcellular localization according to COMPARTMENTS enrichment analysis of all proteins enriched in the BMIF at diagnosis or after induction therapy. **(D)** Comparison of the metabolic enzymes that were enriched in the either leukemic or post-induction BMIF.

Principal Component Analyses (PCA) of quantified proteins showed that the BMIF proteome was clearly distinct from the PBP proteome leading to clear separation based on the first two principal components alone (Figure 1D). Of note, the principal components of some individual samples in the proteomics data deviated from the experimental group mean and aligned more with other experimental groups. However, we examined the available clinical data and were not able to identify any clinical or experimental confounding factors explaining this effect. The metabolite profile of BMIF and PBP also appeared distinct (Figure 1E), while the lipid (Figure 1F) and termini (Figure 1G) profiles do not clearly delineate the two compartments. In contrast to the compartment separation, all analyte groups appear to be remodeled following chemotherapy as seen by separation of diagnostic and post induction specimens based on the first two principal components in any of the data sets. This high-level assessment demonstrated a clear proteome and metabolome difference between compartments and through treatment and suggested that lipid composition and protein processing primarily changed through treatment. In the following we investigated the relationship between analyte groups and characterized specific processes in greater detail.

To integrate protein, protein processing, metabolite and lipid dynamics we trained a UMAP model using all analytes that showed significant changes between any of the conditions. We then mapped the remaining features using this model and used c-means clustering to assign each feature to one of nine clusters (Figure 2A). Each cluster contained analytes from all four analyte classes confirming that the normalization and dimensionality reduction primarily retained the biological dynamics rather than differences in methodology (Figure 2B). In line with our initial assessment that lipids change primarily between timepoints while proteins show strong compartment differences the analyte classes were not evenly distributed across all clusters. Radar plots of the z-scored intensities of all features mapped to a cluster defined 5 clusters representing features enriched in one of the four experimental conditions (clusters 1, 3, 5, 7, and 9), 2 clusters enriched at a timepoint (clusters 2 and 6) and 2 clusters enriched in a compartment (clusters 4 and 8) (Figure 2C).

### 2.2. The BMIF is differentiated from the PBP by proteins localized to extracellular vesicles and exosomes

To explore the differences between the compartments, we turned to differential analysis. At both timepoints, the BMIF showed a larger proportion of enriched proteins. At diagnosis, we found 87 proteins enriched in the BMIF against 4 proteins that were enriched in the PBP (Figure3A); whereas there were 98 proteins that were enriched in the BMIF after induction therapy, compared to 9 proteins that were enriched in the PBP (Figure 3B). In total, 43 proteins were consistently enriched in the BMIF over PBP. COMPARTMENTS enrichment analysis of the BMIF-enriched— either at diagnosis or post-induction— proteins suggested that most of them can be found in extracellular vesicles and exosomes (Figure 3C). In addition, many of the proteins enriched and localized in extracellular structures were enzymes involved in key metabolic processes including glycolysis, lactate production, and glutathione metabolism (Figure 3D).

Hemoglobin subunits also had higher intensities in the BMIF compared to the PBP (Figure 4A), and were uniquely enriched in the BMIF only after induction therapy (Figure 3C). To further investigate this observation, bone marrow biopsies taken at the same time of the bone marrow aspirates were evaluated. Hematoxylin and eosin (H&E) staining showed the expected high number of blast cells at diagnosis diminishing most other cell types. After induction, chemotherapy blasts are eliminated and hematopoiesis increased (Supplementary Figure 1). In addition to an increase of erythrocytes we also noticed an increase of diffuse brown pigmentation resembling hemosiderin deposition (Figure 4B). Quantification of the hemosiderin in bone marrow biopsies (Figure 4C) correlated with hemoglobin levels in BMIF (Figure 4D). Overall, this provided a strong rationale that the observed hemoglobin in the proteomics data originated from *in vivo* hemolysis and demonstrated how the high rate of hematopoiesis in the bone marrow post chemotherapy affects not only the cellular but also the proteome composition.

**Figure 4.**
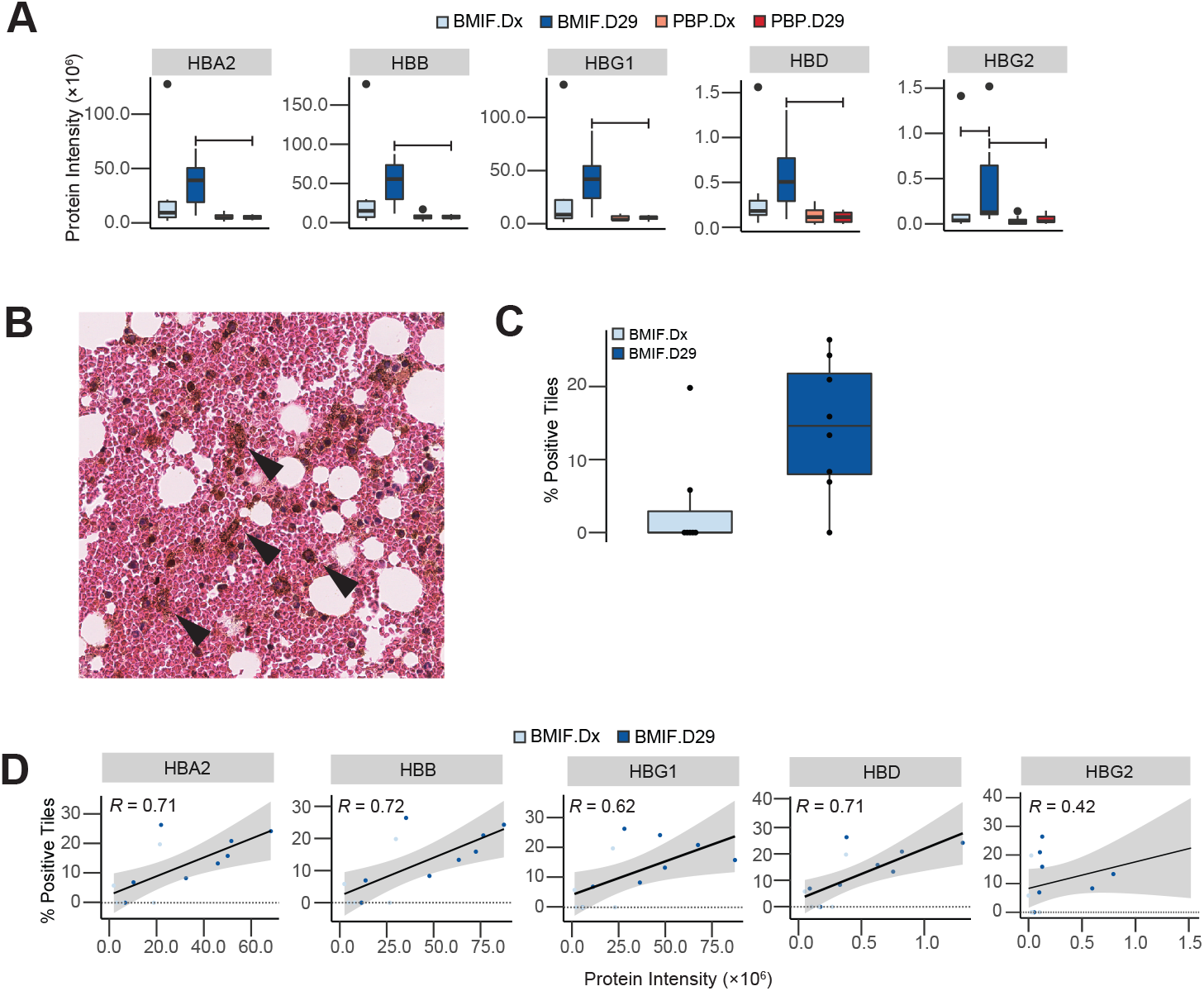
BMIF proteome and bone marrow histology show hemolysis post-induction therapy. **(A)** Intensity of hemoglobin subunits in experimental groups. **(B)** Representative image of hematoxylin and eosin staining (H&E) showing hemosiderin like deposition (arrow) in BMIF. **(C)** Comparison of the hemosiderin-positive tile counts in H&E images of leukemic and post-induction BMIF. **(D)** Correlation between total hemosiderin-positive tile counts and hemoglobin subunit intensities.

### 2.3. Lipid homeostasis is restored after chemotherapy

UMAP dimensionality reduction followed by clustering mapped apolipoproteins in cluster 6 (low PBP.Dx, medium BMIF.Dx, high PBP.D29 & high BMIF.D29), thus suggesting changes in lipidomic profiles during induction chemotherapy (Figure 5A). Of the 14 apolipoproteins detected, 12 had higher abundance after induction therapy in either compartment 10 of which were significantly altered in the PBP compartment (Figure 5B). Coinciding with this were significant increases in the abundance of lipid-soluble vitamin transporters– namely, proteins that functioned in the transport of lipid-soluble vitamins, such as RBP4 (retinol), TTR (retinol), and AFM (Vitamin E) (Figure 5C). In addition, PCSK9, a critical player in lipid homeostasis mediating the degradation of low-density lipoprotein receptors (LDLR), was also significantly elevated after chemotherapy (Figure 5C).

**Figure 5.**
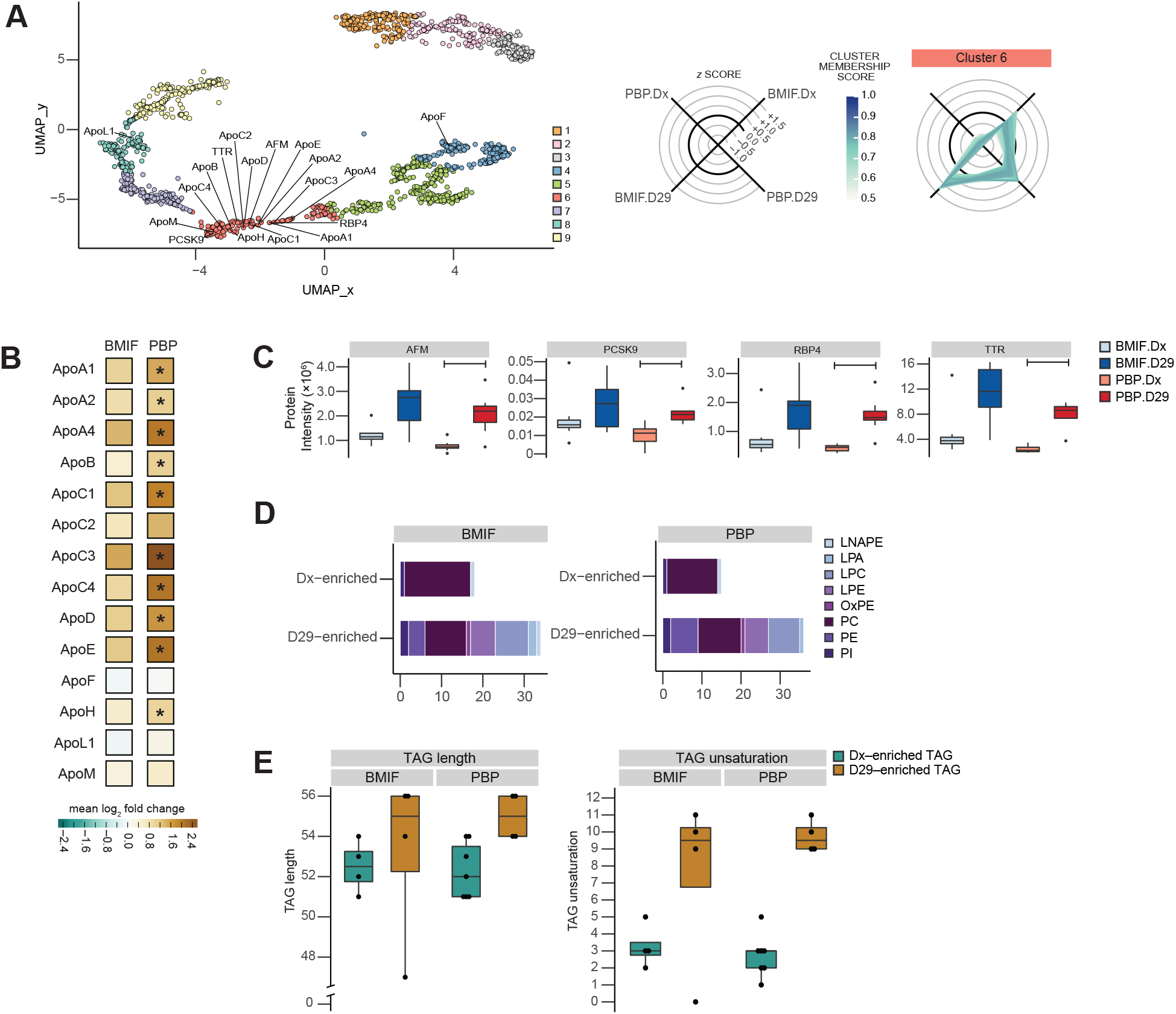
Lipid-binding proteins upregulated after induction therapy and changes in lipidomic profile. **(A)** UMAP of all analytes with apolipoproteins detected in the proteomics data highlighted. **(B)** Fold changes between detected apolipoproteins, with significantly increased (limma, adjusted p-value ≤ 0.05) apolipoproteins from diagnosis to post-induction indicated by (*). **(C)** Protein intensity of lipid binding proteins that were significantly increased at post-induction in PBP differential analysis. Bars indicate significant difference (limma, adjusted p-value ≤ 0.05) between the experimental conditions. **(D)** Classification of D29-en-riched glycerophospholipids. (E) Comparison of the length and unsaturation levels of the triacylglycerols (TAGs) enriched at either diagnosis and post-induction.

To further characterize the lipidomic changes in ALL patients through chemotherapy, we inspected the lipid classes. The glycerophospholipid profile at diagnosis was significantly enriched for phosphatidylcholines (PC) while other sub-classes are significantly de-riched constituting a reduced complexity (Supplementary Figure 4A, Supplementary Figure 4B). After induction therapy, most glycerophospholipids were elevated restoring a more diverse lipid profile in both compartments (Figure 5D). Among the glycerophospholipids that saw bigger increases in the post-induction samples were lysophosphatidylcholines (LPC; from 0% in either Dx samples to 24% and 22% in BMIF.D29 and PB.D29 samples, respectively).

Aside from glycerophospholipids, there were also changes in the glycerolipids; specifically, lipoprotein-associated triacylglycerols. In the PBP compartment, the majority of glycerolipids have significantly higher abundance at diagnosis (Supplemental Figure 4B). Upon additional examination, significantly altered triacylglycerols (TAGs) in either timepoint possessed different properties. After induction therapy, the significant TAGs typically had higher levels of unsaturation and longer length, whereas those that were significantly higher in diagnostic samples were shorter and more saturated (Figure Figure 5E).

### 2.4. Metabolic changes and reduction in proteolytic complement activation form an immunosuppressive microenvironment post chemotherapy

The innate immune system is known to be deregulated in ALL [18]. The protease driven deregulation of the complement system in particular has been associated with pro- and anti-tumorigenic effects in the leukemic microenvironment [19,20]. This established concept was captured in the UMAP representation of our data and confirmed by subsequent clustering analysis. Here, the complement proteins and fragments were split between high-Dx and high-D29 clusters (Figure 6A).

**Figure 6.**
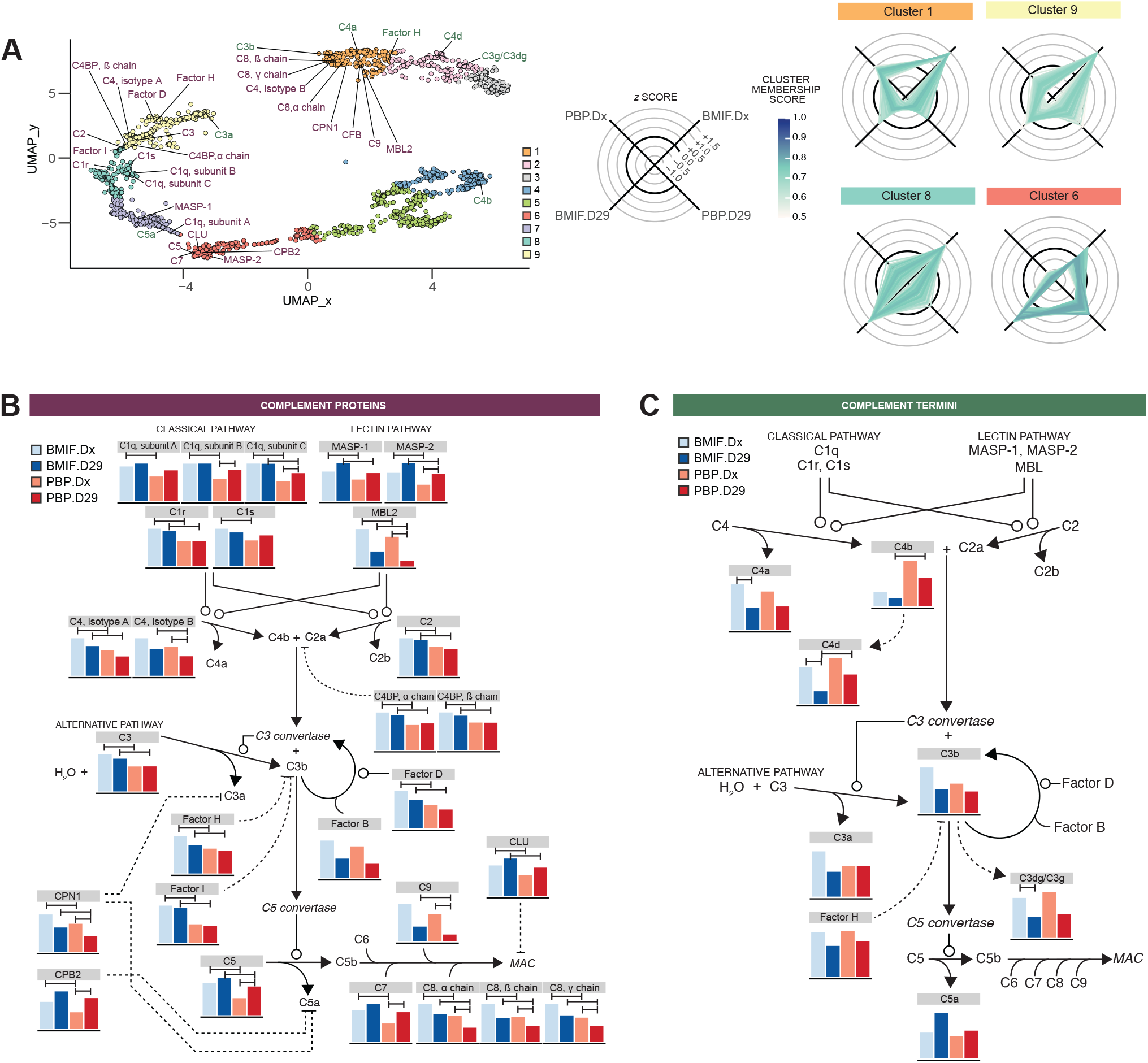
Complement system changes. **(A)** UMAP of all analytes with complement proteins and termini detected in the proteomics and terminomics data, respectively, highlighted. Z-scaled intensity of detected complement **(B)** proteins and **(C)** termini mapped to the complement pathway. Bars indicate significant difference (limma, adjusted p-value ≤ 0.05) between the experimental conditions.

To elucidate the activity of the complement system in the context of the pediatric ALL microenvironment we quantitatively mapped the complement pathway at the protein level and assessed the activation state of pathway segments using quantitation of the complement fragment specific N termini. In total, we mapped 27 proteins (Figure 6B) and 8 termini (Figure 6C) on the complement pathway. In general, complement proteins were significantly more abundant in BMIF than the PBP. Overall, the abundance of complement pathway proteins was stable or moderately reduced following chemotherapy. This reduction was more consistent and pronounced in the PBP. Here in particular MBL2, an activator of the lectin pathway, and components of the MAC complex showed significant changes (Figure 6B).

In the BMIF complement protein abundance changed only moderately, while their proteolytic processing and abundance of the resulting N termini / fragments changed significantly. Following chemotherapy 2 fragments generated from C4 protein (C4a, C4d) and C3 protein (C3g or C3dg) were not only significantly lower than during overt leukemia but also lower than in the post-chemotherapy PBP (Figure 6C).

In addition to the signature of reduced complement activation we found a significant decline of immune system components in the PBP, including LRG, ICAM1 and MARCO (Supplementary Figures 5A, 6A). In the BMIF, the immune proteins were not robustly identified and its alteration, therefore, could not be statistically defined. However, we observed significantly reduced levels of extracellular adenosine that provided further evidence of immunosuppression post-induction therapy (Supplementary Figures 6B). Extracellular adenosine, a metabolite of ATP, has profound effects on the microenvironment. One proposed mechanism by which extracellular adenosine dampens immune activity is via the A_2A_ receptor. This results in downstream cAMP signaling with an associated induction of SOCS-3 protein and suppression of JAK-STAT pathway [21]. To determine the relevance of this mechanism in the context of the post leukemic BMIF, we evaluated levels of Gp130. Gp130 is part of the initialization of JAK-STAT pathway, but it is also integral to the negative feedback loop via SOCS-3 proteins. Furthermore, the soluble form of gp130 has been previously detected in plasma, and its capability at countering anti-apoptotic effects of IL-6R-mediated signaling has been previously explored [22]. We found that gp130 was detected almost exclusively in post-induction samples (Supplementary Figure 6C). Post-induction it was significantly more abundant in the BMIF compared to the PBP. Together, our findings provide evidence for an immunosuppressed microenvironment, in particular in the bone marrow, following induction therapy.

## 3. DISCUSSION

In the present study, we hypothesized that the BMIF and PBP microenvironments are molecularly distinct. More specifically, we aimed to investigate the fluid portion of the microenvironment, which has been under-explored in previous studies. In addition, we also explored how these compartments change in the context of pediatric B-ALL over the course of chemotherapy. To do this, we examined liquid patient biopsies from two compartments (bone marrow interstitial fluid and peripheral blood plasma) and taken at two timepoints (diagnosis and post-induction).

In our analyses, the BMIF was enriched for proteins that are known to be localized in extracellular exosomes and vesicles (EVs). These organelles have previously been shown to play important roles in malignant and normal settings, matching their enrichment in the BMIF regardless of timepoint [23–25]. However, to the best of our knowledge, the current study is the first to report the enrichment of EV-localized proteins, including key metabolic enzymes, in the BMIF compared to the PBP. There is a growing interest in the clinical significance of EVs and our work demonstrates clear potential in the context of pediatric ALL but also emphasizes the need for further studies to determine if modulation of EVs could constitute a viable diagnostic or therapeutic approach [26].

Together, our proteomic and lipidomic analyses demonstrated significant increase in lipid-binding proteins—including apolipoproteins—and changes in lipid composition. Our findings are in line with previous studies that have demonstrated lipidomic changes in ALL patients after chemotherapy [27,28]. The apolipoproteins that we found significantly changed, namely Apo-B, Apo-E, Apo-C1, and Apo-C3, are known to be associated with atherogenic lipoproteins. Coupled with a significant increase in PCSK9 levels in the peripheral blood after induction therapy, this confirms studies that have examined the development of metabolic syndromes, including dyslipidemia and cardiovascular diseases in ALL survivors [29–31].

After induction therapy, both the BMIF and the PBP were altered towards a post-apoptotic and immunosuppressive microenvironment. In the BMIF, this included the hemolytic aftermath signified by increased levels of hemoglobin subunits and hemosiderin deposition, and the elevated levels of extracellular adenosine. In the PBP, we observed significant dampening of the complement system and other immune-related proteins. Our findings reinforced previous studies that have sought to determine the effects of induction therapy agents. For instance, MBL2 levels observed post-induction could be attributed to the effects of asparaginase treatment [32]. The mostly exclusive detection of gp130/IL6ST in post-induction samples could be linked to glucocorticoids such as dexamethasone or prednisolone [33]. The presence of extracellular adenosine has been previously suggested to be a result of the drug methotrexate [34,35].

While the cellular immune response is reinforced by increased hematopoiesis following induction chemotherapy our data confirms substantial reduction in the humoral innate immune response in line with increased susceptibility to infection[36,37]. Importantly, elements of this post-apoptotic immunosuppressed humoral microenvironment, such as elevated adenosine, may play an essential role in the regeneration of the hematopoietic progenitor cell pool [38]. Thus, the co-occurrence of immunosuppression and restoration of hematopoiesis following induction therapy warrants additional exploration. Controlled modulation of the post-chemotherapy microenvironment, e.g., to tune the balance between necessary immunosuppression and vulnerability to infection, could constitute an effective avenue to reduce chemotherapeutic side effects.

## 4. CONCLUSION

Through our multi-omic analysis, we found that the BMIF and PBP provided different contributions to the tumourigenesis of ALL and underwent different changes through chemotherapy. Particularly, we found that the BMIF harboured elevated levels of extracellular vesicle associated proteins that could be instrumental in cross-talk between leukemic cells and cells in the microenvironment. Lipid homeostasis was restored following chemotherapy. Immune regulatory proteins that were highly abundant in the leukemic PBP decreased over the course of chemotherapy while the BMIF switches in favour of a post-apoptotic and immunosuppressive microenvironment. Our study further suggests that monitoring of select activated complement fragments rather than the endpoints may serve as more sensitive biomarkers of the complement status.

## 5. ACKNOWLEDGMENTS

This work was partially supported by grants from the Michael Cuccione Foundation, the BC Children’s Hospital Foundation (to P.F.L.), and the Canadian Institutes of Health Research (PJT-169190 to P.F.L.). L.N. was supported by a fellowship from the Michael Cuccione Childhood Cancer Research Program. P.F.L. was supported by the Canada Research Chairs program and the Michael Smith Foundation for Health Research Scholar program.

We thank Dr. Seth Parker and Enes Kemal Ergin for assistance and consultation. We are also grateful for the patients and families who participated and made this study possible, as well as the BC Children’s Hospital nurses, physicians, and staff at the Biobank for their collective effort in acquiring and maintaining specimens.

## 6. METHODS

### 6.1. Patient samples

Acute Lymphoblastic Leukemia (ALL) patient samples and non-ALL patient control samples were obtained from the Biobank at the BC Children’s Hospital (BCCH) following informed patient consent and approval by the University of British Columbia’s and Women’s Research Ethics Board (H15-01994) in agreement with the Declaration of Helsinki. Patient bone marrow interstitial fluid (BMIF) and matching peripheral blood plasma (PBP) were collected at the time of diagnosis (Dx, pre-treatment) and 29 days after initiation of the treatment at the end of the induction phase of chemotherapy (D29, post-treatment).

### 6.2. Proteomic and terminomic analyses

#### 6.2.1. General sample preparation for proteomic and terminomic samples

A 5 µL aliquot from each sample was transferred into a 1.5 mL Protein LoBind tube (Eppendorf) that contained 15 µL lysis buffer and 1 µL 25U/µL Benzonase (Millipore). The lysis buffer consisted of 1% sodium dodecyl sulphate (SDS, Fisher BioReagents) and 1 × Halt^™^ Protease Inhibitor Cocktail (Thermo Scientific) in 50mM HEPES, pH 8.0 (Sigma). The lysate was incubated at 37°C for 30 minutes. Samples were then heated at 95°C for 5 minutes which was immediately followed by a 5-minute incubation on ice. Protein concentration was estimated using Bicinchoninic acid (BCA) assay.

Samples were reduced and alkylated using 2 µL of 100mM Dithiothreitol (DTT, 10mM final concentration) and 2 µL of 500mM Chloroacetamide (CAA, 50mM final concentration), respectively. Samples were incubated at 37°C for 30 minutes during reduction, while alkylation was done at room temperature, in the dark, for 30 minutes. To quench the alkylation reaction, 2 µL of 1M DTT was added. Reduced and alkylated samples were flash-frozen in liquid nitrogen and stored at −80°C.

#### 6.2.2. Single-pot, solid-phase-enhanced sample preparation (SP3) for proteomic analysis

Samples were processed for proteomic analyses in two batches, with each batch having 18 samples each. Samples were selected randomly and were collected from −80°C storage. After the samples thawed, a 5.2 µL (equivalent to 20% of the processed sample, or 1 µL of undiluted BMIF/PBP) aliquot was transferred into a 1.5 mL Protein LoBind tube and was diluted with 4.8 µL 50mM HEPES, pH 8.0. Prepared SP3 beads (10 µL hydrophobic and 10 µL hydrophilic) were added to the lysate and bead binding was initiated through the addition of 80 µL non-denatured Ethanol (EtOH). After 18 minutes of incubation at room temperature, with end-over-end mixing, the SP3 beads were magnetically isolated and the supernatant was removed. SP3 beads were washed three times with 200 µL 90% EtOH. After removing the last wash, the SP3 beads were briefly centrifuged to remove excess EtOH. SP3 beads were re-suspended in 200mM HEPES, pH 8.0 and incubated with trypsin at 1:75 protease:protein (w/w) ratio. Samples were incubated overnight at 37°C with constant agitation.

After digestion, SP3 beads were magnetically isolated and the digested peptides were desalted for LC-MS/MS analysis using BioPureSPN MINI column (The Nest Group Inc.) according to the manufacturer’s instructions. Briefly, the columns were conditioned by sequentially loading 200 µL Methanol (MeOH), 200 µL Buffer B, and 200 µL Buffer A. Buffer A consisted of 0.1% Trifluoroacetic acid (TFA) in water and Buffer B consisted of 0.1% Formic acid (FA) in 60% Acetonitrile (ACN). Samples were prepared for the column by topping it up to final volume of 100 µL using Buffer A and acidified (pH=2.0– 3.0) with 4 µL 10% TFA. Samples were loaded through the column and the flow through was collected. The collected flow through was re-loaded into the column. The columns were washed with three times with 200 µL Buffer A. Peptides were eluted from the column using a total of 190 µL Buffer B. Acetonitrile in the collected peptides was removed by speed-vac.

#### 6.2.3. Automated High-efficiency Undecanal-based N Termini EnRichment (HUNTER) for terminomic analysis

Samples were processed for terminomic analyses in four batches, with each batch having 9 samples each. Samples were selected randomly and were collected from –80°C storage. From the remaining 80% of the prepared sample, an aliquot equivalent to approximately 100 µg of starting material was transferred into a 1.5 mL Protein Lobind microcentrifuge tube. Samples were topped up to a final volume of 20.8 µL using 50mM HEPES, pH 8.0 before it was transferred to the twin.tec PCR plate 96 (semi-skirted; Eppendorf). Samples were processed with the program and settings previously established for the Automated HUNTER workflow [14] using epBlue Studio (version 40.4.0.38). Samples were processed on an epMotion M5073 automated liquid handling system (Eppendorf) controlled by an EasyCon tablet (Eppendorf). In addition, the M5073 automated liquid handling system was also configured with the following accessories: dispensing tool TS50 (1.0 – 50 µL) and TS1000 (40 – 1000 µL), epT.I.P.S. Motion racks (1.0 – 50 µL and 40 – 1000 µL), epMotion gripper, Thermoadapter for 96-PCR plate (skirted), Alpaqua Magnum FLX 96 magnet plate, Eppendorf rack for 24 × safe lock.

Sample processing took two days, with the first day consisting of SP3 bead binding, dimethylation, quenching, SP3 bead re-binding, and trypsin digestion. Dimethylation was done by adding 2M Formaldehyde (CH_2_O) and Sodium cyanoborohydride (NaBH_3_CN) until a final concentration of 40mM and 20mM, respectively. Dimethylation was done at room temperature for 1 hour with the sample plate covered with Thermal Adhesive Sealing Film (Diamed). Trypsin digestion was done at a 1:70 protease:protein (w/w) ratio in 30 µL 200mM HEPES, pH 8.0. Sample plate was covered with the Thermal Adhesive Sealing Film and was incubated at 37°C overnight. On the following day, the sample plate was briefly centrifuged to collect condensation in the film. Undecanal labeling was initiated by adding 5 µL Undecanal and NaBH_3_CN was added. Sample plates were covered with Thermal Adhesive Sealing Film and the labeling reaction was allowed to proceed at 37°C for 1 hour.

Samples were collected from the plate and transferred into 1.5 mL Protein LoBind microcentrifuge tubes. The removal of excess Undecanal and depletion of undecanal-labeled internal peptides was done using Sep-Pak. Briefly, tC18 Sep-Pak (Waters) columns were conditioned by sequentially loading 700 µL Methanol and 700 µL 0.1% TFA in 40% EtOH. Samples were prepared for loading by mixing it with 700 µL 0.1% TFA in 40% EtOH and acidifying it (pH=2.0–3.0) with 7 µL 10% TFA. Samples were loaded and the flow through was collected. The flow through contains peptides that were not excluded through undecanal label. EtOH in the flow through is removed through speed vac.

Samples were processed through a second round of clean up using STaGE tips. STaGE tips were assembled with four circular Solid Phase Extraction (SPE) C18 disks (Empore) that were created by using a flat-end needle as a puncher. The C18 disks were stacked in a P200 pipette tips using a straightened paper clip. The assembled STaGE tips were conditioned by loading 80 µL Methanol followed by 80 µL 0.1% TFA in water. Samples were topped up with 0.1% TFA in water to a final volume of 80 µL, if necessary. Samples were loaded through the STaGE tip and the flow through was collected and re-loaded. STaGE tips were washed two times with 0.1% TFA in 2% ACN. Peptides were eluted from the STaGE tip using 80 µL 0.1% FA in 40% EtOH. EtOH was removed from the samples using the speed vac.

#### 6.2.4. Sample pool fractionations

Sample pools were prepared by taking 2 µL aliquots per sample. The sample pool for proteomic samples was depleted by using Top 2 Abundant Protein Depletion Spin Columns (Pierce) then subsequently Top 12 Abundant Protein Depletion Spin Columns (Pierce) according to manufacturer’s protocol. After protein depletion, sample pools were processed by SP3 as described previously. For terminomics, the sample pool was not depleted but was processed by Automated HUNTER as described previously. Offline high-pH reversed phase fractionation was performed on the depleted proteomic and non-depleted terminomic sample pools. An Agilent 1100 HPLC system equipped with a diode array detector (254, 260, and 280 nm) and a Kinetic EVO C18 column (Phenomenex, 2.1 mm 150 mm, 1.7µm core shell, 100 pore size) was used. Mobile phase A was 10 mM Ammonium bicarbonate, pH 8.0 and mobile phase B was Acetonitrile. The gradient was 3% to 35% B over 60 min. Samples were run at a flow rate of 0.2 ml per min. Fractions were collected every minute across the elution window for a total of 48 fractions. Fractions collected from the depleted proteomic sample pool were concatenated to a final set of 12. Fractions collected from non-depleted terminomic sample pools were concatenated to a final set of 6. Samples were dried in a SpeedVac centrifuge and re-suspended in 0.1% FA in water prior to DDA mass spectrometry analysis.

#### 6.2.5. LC-MS/MS analysis of proteomic and terminomic samples

Proteomic and terminomic analyses (including fractions) were performed on a Q Exactive HF Orbitrap mass spectrometer coupled to an Easy-nLC 1200 liquid chromatography system (Thermo Scientific). Mobile phase A was 2% ACN in 0.1% FA in water and Mobile phase B was 95% ACN in 0.1% FA in water. A 20 cm-long PicoFrit^™^ (New Objective, 360OD, 75ID, 15µm tip ID) analytical column with a 3 cm-long home-made (Polymicro Technologies, 360OD, 100ID) pre-column was used. Both columns were packed with ReproSil-Pur 120 C18 beads (Dr. Maisch GmbH, 3µm). The analytical column temperature was set to 50°C.

##### 6.2.5.1. Chromatographic gradient of proteomic and terminomic samples

Chromatographic gradient was set as follows: 0 min: 3% B; 2 min: 8% B; 42 min: 27% B; 57 min: 42% B; 60 min: 50% B; 70 min: 50% B. Flow rate was at 0.3 µL/min.

##### 6.2.5.2. Data-independent acquisition (DIA) of proteomic samples and terminomic samples

A full scan MS spectrum (300–1650 *m/z*) was collected with a resolution of 120,000. Maximum injection time was 60 ms and AGC target value was 3e6. DIA segment spectra were acquired with 24-variable window format with a resolution of 30,000. AGC target value was 3e6. Maximum injection time was set to ‘auto’. The stepped collision energy was set to 25.5, 27.0, 30.0.

##### 6.2.5.3. Data-dependent acquisition (DDA) of depleted proteomic fractions

For data-dependent acquisition (DDA) of depleted proteomic fractions (i.e., prepared via offline high-pH fractionation), a full scan MS spectrum (350–1650 *m/z*) was collected with a resolution of 120,000. Maximum injection time was 30 ms and an AGC target value of 2e5 with an isolation window of 1.4 *m/z*. The top 12 precursors were selected for fragmentation. Normalized collision energy was set to 28. Dynamic exclusion duration was set to 15 s. Charge state exclusion was set to ignore unassigned, 1, 5 – 8, and greater.

##### 6.2.5.4. DDA of depleted non-depleted terminomic fractions

For DDA of non-depleted terminomic fractions (i.e., prepared via offline high-pH fractionation), a full scan MS spectrum (350–1600 *m/z*) was collected with a resolution of 120,000. Maximum injection time was 246 ms and an AGC target value of 1e6 with an isolation window of 1.4 m/z at Orbitrap cell. The top 12 precursors were selected. Normalized collision energy was set to 28. Dynamic exclusion duration was set to 15 s. Charge state exclusion was set to ignore unassigned, 1, 5 – 8, and greater.

#### 6.2.6. Spectral library preparation

Proteomic spectral library was prepared using Spectronaut (version 13.9.191106.43655). The proteomic spectral library was comprised of the individual DIA samples, and fractionated DDA samples. Enzyme and digestion type was set to Specific and Trypsin/P, respectively. Maximum and minimum peptide length were set to 7 and 52 amino acids, respectively. Missed cleavage was set to 2. Acetyl (Protein N-term) and Oxidation (M) were set as variable modifications; Carbamidomethyl (C) was set as fixed modifications. The proteomic spectral library yielded 759 protein groups.

The terminomic spectral library was comprised of the fractionated DDA samples supplemented with additional DDA runs from a previous study that also utilized HUNTER on B-ALL patient plasma samples [14]. To prepare the spectral library, DDA runs were initially processed using MaxQuant (version 1.6.10). Enzyme and digestion type was set to Semi-specific free N-terminus and Trypsin/P, respectively. Minimum peptide length was set to 7. Oxidation (M), Dimethyl (N-term), and Acetyl (N-term) were set as variable modifcations; Carbamidomethyl (C) and Dimethyl (K) were set as fixed modification. After the MaxQuant search, a spectral library was prepared on Spectronaut (version 14.3.200701). The spectral library from the fractionated DDA samples yielded 196 protein groups and 1,266 modified peptides from 1,619 precursors, while the supplemental library had 218 protein groups and 1,601 modified peptides from 2,065 precursors.

#### 6.2.7. Data extraction

Proteomics data was prepared using Spectronaut (version 13.9.191106.43655) using the prepared spectral library, described above, and the default BGS Factory Settings. Terminomics data was prepared using Spectronaut (version 13.9.191106.43655) using the prepared spectral libraries, described above, and the following modifications to the default BGS Factory Settings: Single Hit Modification was set to Modified Sequence in the Identification tab; Minor (peptide) Group was set to by Modified Sequence, and the Minimum Major Group Top N was set to 2 under the Quantification tab.

### 6.3. Metabolomic and lipidomic analyses

#### 6.3.1. Dual Phase Extraction

When available, 50 µL aliquot samples were quenched by adding 270 µL ice cold methanol and placed in −20 °C freezer overnight to sediment the precipitated proteins. Then 900 µL methyl tert-butyl ether (MTBE) was mixed with the extracted solution and vortexed for 30 min on an open-air microplate shaker. 265 µL H_2_O was then added into the solution followed by centrifugation (14000 rpm, 4°C, 1 min) to accelerate the phase separation. The upper organic phase and lower aqueous phase were separately transported to two 1.5 mL Eppendorf vials and concentrated by speed-vac at 4°C. The concentrated pellets were subsequently reconstituted with (ACN:IPA=1:1, v:v) for the lipids from organic phase and (ACN:H_2_O=1:1, v:v) for the metabolites from aqueous phase. One method control (MC) was prepared using the same procedure but with no real sample in the vial. One quality control (QC) was prepared by pooling 5 µL from each of the solutions. QC was injected between every 4 sample injections to assure the instrument was stable and the sample injection volume was optimized.

#### 6.3.2. LC-MS Analysis

Metabolomic and lipidomic LC-MS analyses were performed on Bruker Impact II(tm) UHR-QqTOF (Ultra-High Resolution Qq-Time-Of-Flight) mass spectrometer coupled with the Agilent 1290 Infinity(tm) II LC system. 2 µL sample solutions of each replicate. A Waters ACQUITY UPLC BEH C18 Column (130Å, 1.7 µm, 1.0 mm X 100 mm) was used for lipidomics in RP(+) and (-). The mobile phase A was 60% ACN 40% H_2_O (2 mM NH_4_Ac in positive mode, 5 mM NH_4_Ac in negative mode); mobile phase B was 90% IPA 10% ACN (2 mM NH_4_Ac in positive mode, 5 mM NH_4_Ac in negative mode). The chromatographic gradient was set as follows: 0 min: 95% A; 8 min: 60% A; 14 min: 30% A; 20 min: 5% A; 23 min: 5% A. Post-gradient equilibrium time was 9 min at 95% A. Flow rate was 0.1 mL/min. A Millipore ZIC-pHILIC column (200Å, 5 µm particle size, 2.1 × 150 mm) was used for metabolomics in HILIC(+) and (-). Mobile phase A was 95% H_2_O and 5% ACN (pH=4.8 in positive mode, pH=9.8 in negative mode); mobile phase B was 5% H_2_O and 95% ACN. The chromatographic gradient was set as follows: 0 min, 5% A; 20 min, 80% A; 25 min, 95% A. Post-gradient equilibrium time was 19 min at 5% A. Flow rate was 0.15 mL/min during the gradient, and 0.2 mL/min during the post-gradient equilibrium. The mass spectrometer was operated in Auto MS/MS. The ionization source capillary voltage was set to 4.5 kV in positive scanning mode and −3.5 kV in negative scanning mode. The nebulizer gas pressure was set to 1.0 bar. The dry gas temperature was set to 220 °C. The collision energy for MS/MS was set to 7 eV. The data acquisition was performed in a range of 50-1200 m/z at a frequency of 8 Hz.

#### 6.3.3. Data extraction

Bruker DataAnalysis software was used to calibrate the spectra using NaFA as the internal reference. Then the raw data files with format of ‘.d’ were converted to format of ‘.abf’ using AbfConverter. The converted files were then uploaded onto MS-DIAL for both metabolite and lipid feature extraction and alignment. The metabolites were annotated by MS^2^ matching against the publicly available MS^2^ spectral database provided by MS-DIAL (MSMS-Public-Pos-VS11 and MSMS-Public-Neg-VS11). The library contains accurate mass and tandem MS information for annotation. The positive mode library includes the tandem MS information for 29,269 metabolites and the negative mode library includes the tandem MS information for 17,810 metabolites. The tandem MS information was collected from Massbank, ReSpec, GNPS, Fiehn lab, CASMI2016, MetaboBase, and RIKEN PlaSMA. Lipids identification was carried out using LipidBlast available in MS-DIAL [39,40]. The parameters were set as follows: MS1 tolerance was 0.01 Da; MS2 tolerance was 0.05 Da; retention time tolerance was 0.5 min. The alignment result was exported in a ‘.csv’ file.

### 6.4. Data analysis

#### 6.4.1. Data processing

Using the spectral libraries prepared for the terminomics data, we cumulatively identified 1,533 peptides. Initial assessment of the labeling at the N terminus showed that, on average, the 59% of N termini were labeled with dimethyl, whereas 7% were labeled with acetyl, and 34% unlabeled (Supplementary Figure 2). For subsequent analyses, we prepared a deduplicated terminomics data set. Here, peptides that had a label at the N terminus (i.e., dimethyl or acetyl) were merged and considered as one peptide if it had the same N terminal position in the protein. In the deduplicated data set, there was a total of 915 peptides that were labeled (i.e., acetylated or dimethylated) at the N terminus.

Metabolomic and lipidomic data sets acquired from the same liquid chromatography method, albeit using different polarities were merged into one data set (e.g., identifications from HILIC at positive and negative polarities were merged). Entries that had an “Unknown” metabolite or lipid name or did not have MS2 (i.e., designated “w/o MS2”) were excluded from further analysis. Entries that have the same names were collapsed into one entry, with the entry having higher intensities kept.

#### 6.4.2. Normalization and imputation

Normalization was done using the ‘limma’ package in R and the ‘normalizeBetweenArrays’ function. Missing data in all of the data sets were imputed.

#### 6.4.3. Gene Ontology (GO) enrichment analysis

GO enrichment analysis to determine subcellular localization of proteins identified from SP3 and HUNTER were performed using enrichR (mayaanlab.cloud/Enrichr) [41–43] and the COMPARTMENTS[15] database. Outputs from enrichR analysis was plotted using R and the ‘ggplot2’ package.

#### 6.4.4. Principal Component Analysis (PCA)

PCA was done using imputed data sets. Only analytes that we detected in at least 75% of the samples were included for PCA. Data filtering was done using the R and the ‘dplyr’ package. The plots were generated using R and the ‘fviz_pca’ function.

#### 6.4.5. Statistical analyses

Statistical analyses were conducted using R and paired Linear Models for Microarray Data (limma) [44] and imputed data sets. For the statistical analyses, imputed values were assigned with a lower weight (i.e., weight=0.0001) compared to non-imputed values which were assigned with weight=1.0. An analyte was determined to be significant if it has a coefficient ≥ |2|, an adjusted p-value ≤ 0.05, and identified in at least 4 samples within an experimental group. P-value adjustment for multiple testing was done using the p.adjust function and Benjamini-Hochberg method in R. The volcano plots were generated using R and the ‘ggplot2’ package.

#### 6.4.6. Uniform Manifold Approximation and Projection (UMAP)

UMAP was done by merging all of the imputed data sets. Corresponding proteomic and terminomic data were excluded for samples that were not processed for metabolomic and lipidomic analyses (i.e., PBP.Dx for B-ALL #04 and BMIF.Dx for B-ALL #05). After the data sets were merged, the mean intensity of the analytes per experimental group was calculated. The mean intensities were z-score normalized and z-scores were capped at +2 and –2. Analytes that had greater than +2 or less than –2 post-normalization were converted to +2 or −2, respectively. After normalization, the data set was filtered for statistically significant analytes. This data set comprised of statistically significant analytes was used as a training data for UMAP. To prepare the training set, the ‘umap’ function of the ‘umap’ package in R was used with the following settings: n_neighbours=300, min_dist=0.1. To prepare the projection, the complete z-score normalized and capped data set was processed using the ‘predict’ function. For fuzzy-c clustering, we applied the ‘fanny’ function on the projection and defined for 8 clusters to be created. The UMAP (with its various annotations) and radar plots were prepared using the ‘tidyverse’ package in R. In the radar plots, analytes with a cluster membership score of ≥ 0.5 were plotted.

### 6.5. Quantification of hemosiderin in bone marrow cores

Bone marrow cores were stained with hematoxylin and eosin (H&E). Analysis of bone marrow cores were done using QuPath. Once uploaded, a border was drawn around the bone marrow cores. Images were then overlayed with 100 µm tiles. When possible, a minimum of 200 tiles, evenly dispersed throughout the image were assessed for hemosiderin. Results of the hemosiderin assessment were prepared using R and and the ‘tidyverse’ package.

## Supplementary Tables and Figures

**Table 1 | Summary of patient information and clinical data.**

**Figure S1.**
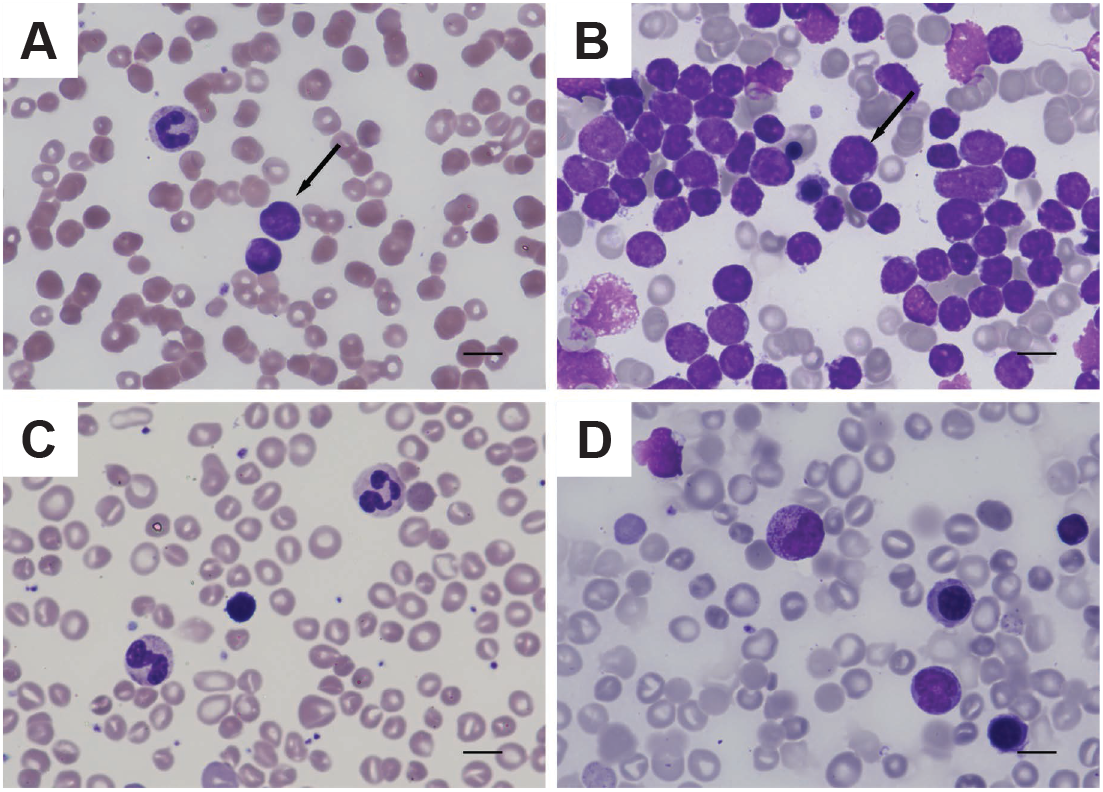
Bone marrow and peripheral blood at diagnosis and after induction therapy. Representative images at diagnosis show blasts (arrows) circulating in the peripheral blood smear **(A)** accompanied with anemia, neutropenia, and thrombocytopenia, while the vast majority of cells in the bone marrow aspirate **(B)** are blasts in a background of reduced trilineage hematopoiesis. Day 29 peripheral blood smear **(C)** shows anisopoikilocytosis with stomatocytes and polychromasia. Day 29 bone marrow aspirate **(D)** shows no definitive blast morphology in a background of increased erythropoiesis and orderly granulopoiesis. All photomicrographs taken at 500x original magnification (oil), scale bar represents 10 microns.

**Figure S2.**
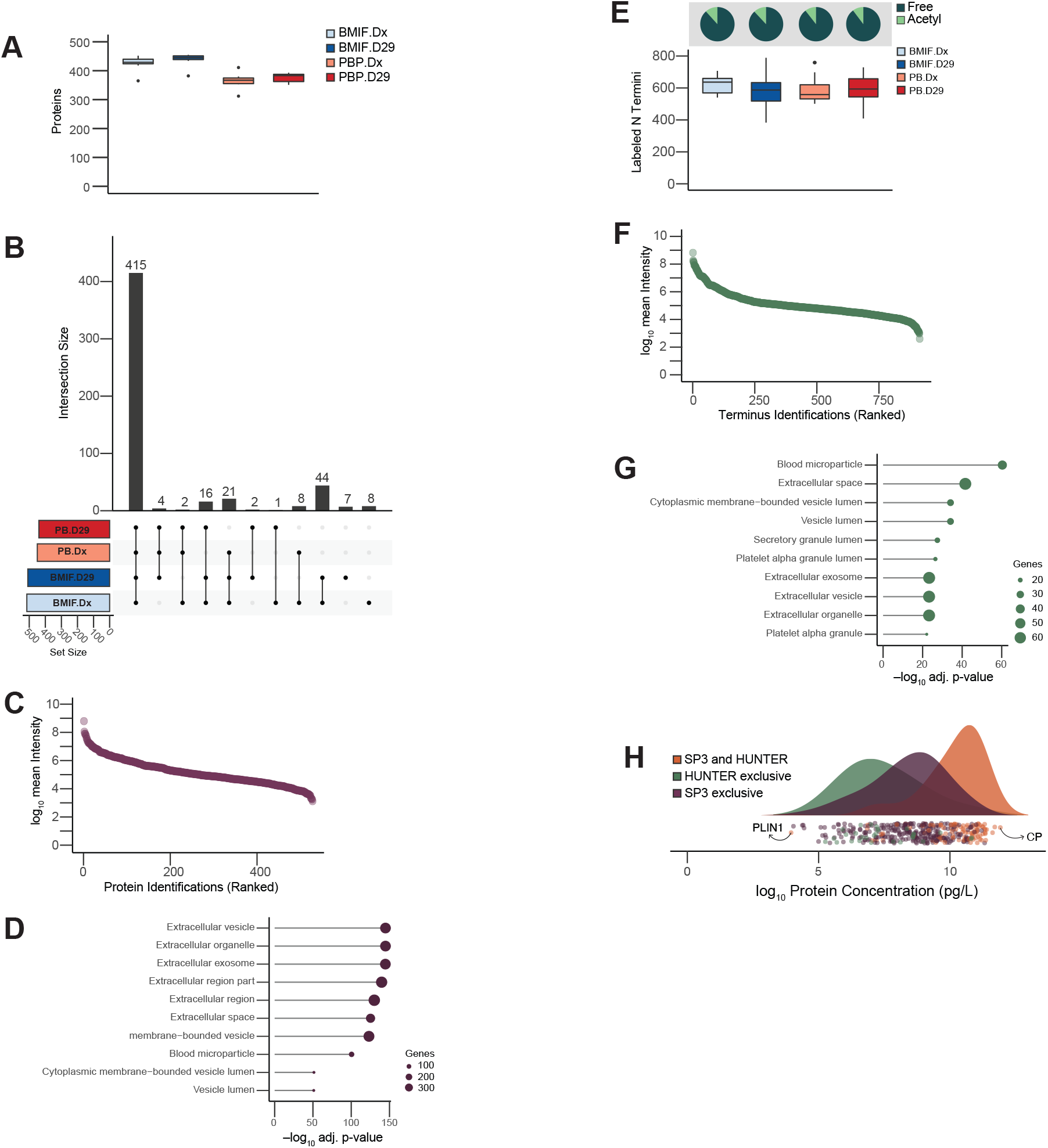
SP3 and HUNTER workflows provide complementary analysis of microenvironment. **(A)** Proteomic identifications per experimental group. **(B)** Upset plot for proteomic identifications. Each bar represents the number shared proteins between the indicated experimental groups **(C)** Mean intensity of proteomic identifications. **(D)** Enriched subcellular localizations in COMPARTMENTS analysis of proteomic identifications. **(E)** Terminomic identifications per experimental group. Pie charts specify the fraction of acetylated and unmodified (free) N termini **(F)** Mean intensity of terminomics identifications. **(G)** Enriched subcellular localizations in COMPARTMENTS analysis of proteins in the terminomic identifications. **(H)** Comparison of the expected plasma concentration of SP3- and HUNTER-detected proteins.

**Figure S3.**
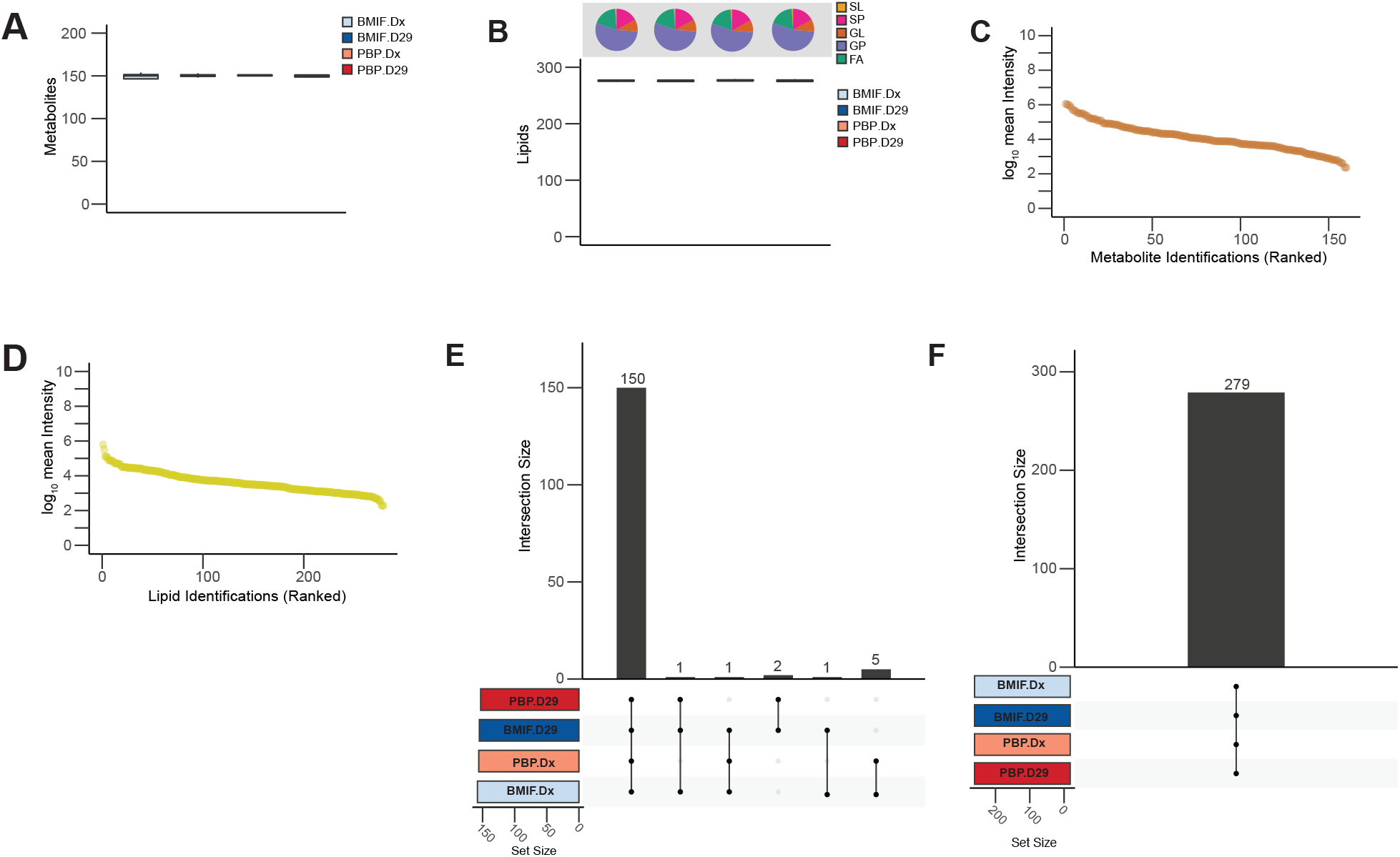
Lipidomic and metabolic identifications were robust. **(A)** Metabolomic identifications per experimental group. **(B)** Lipidomic identifications per experimental group. Pie graphs located at the top represent the proportion of different lipid classes per experimental group (SL: sterol lipids; SP: sphingolipids; GL: glycerolipids; GP: glycerophospholipids; FA: fatty acyls). **(C)** Mean intensity of metabolomic identifications. Mean intensity of lipidomic identifications. **(E)** Upset plot for metabolomic identifications. Each bar represents the number of shared metabolites between the indicated experimental groups. **(F)** Upset plot for lipidomic identifications. Each bar represents the number of shared lipids between the indicated experimental groups.

**Figure S4.**
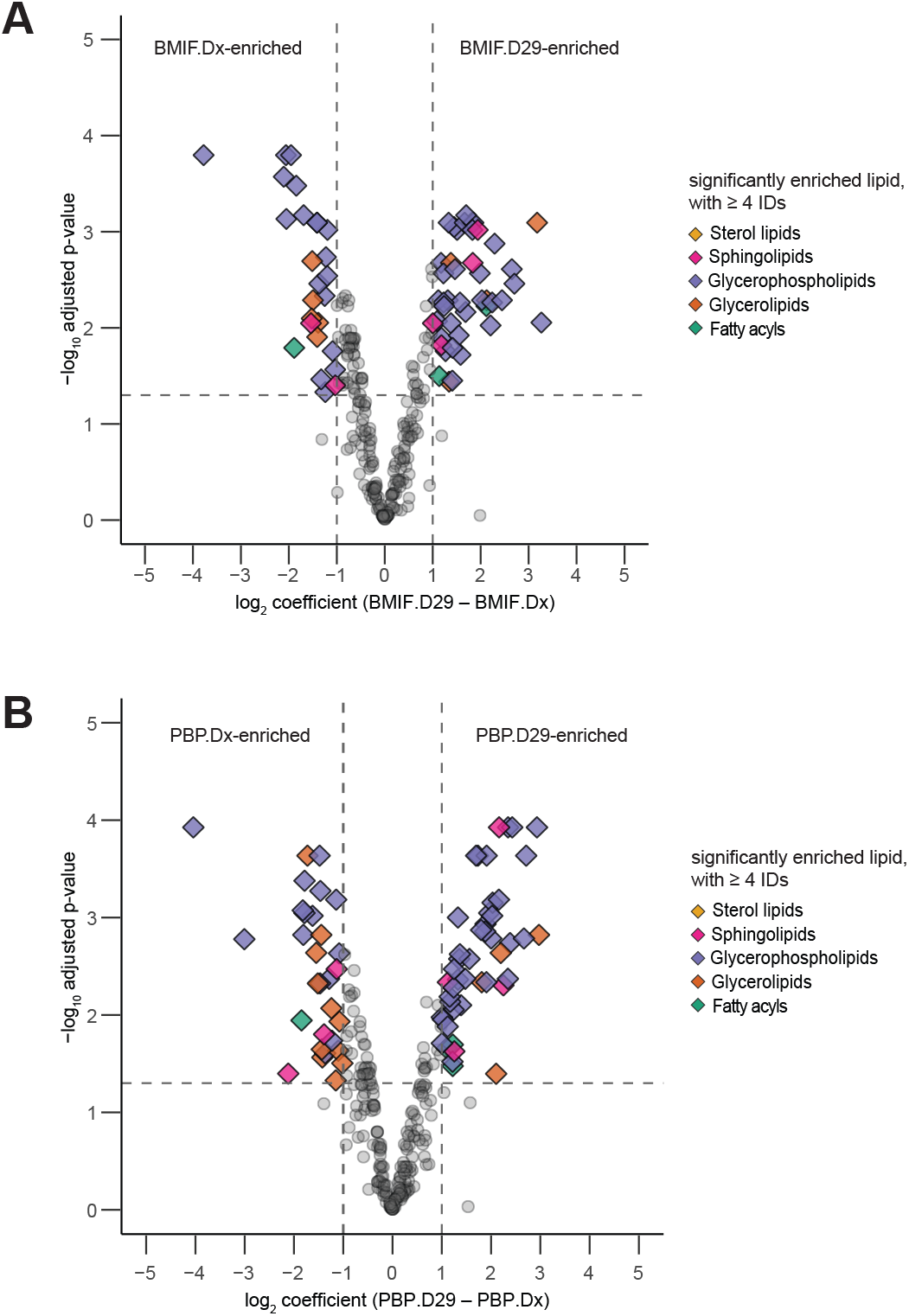
Differential analysis of lipidomics data. **(A)** Temporal changes within BMIF (BMIF.Dx vs BMIF.D29). **(B)** Temporal changes within PB (PBP.Dx vs PBP.D29). Significantly altered lipids are annotated based on lipid class.

**Figure S5.**
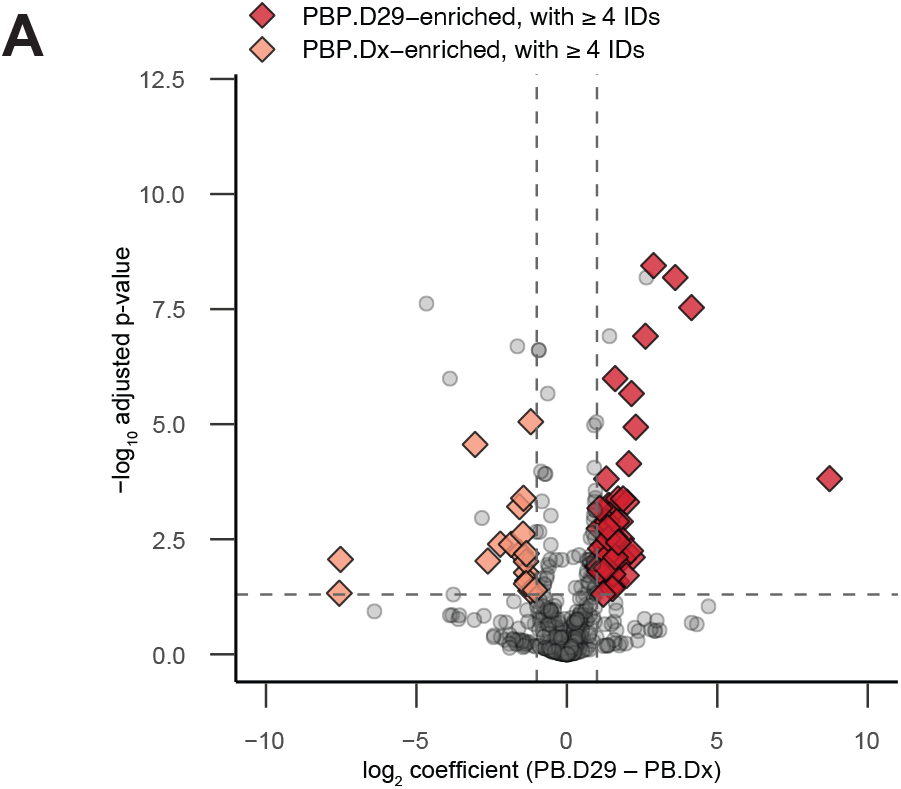
Differential analysis of proteomics data. **(A)** Temporal changes within in PBP (PBP.Dx vs PBP.D29).

**Figure S6.**
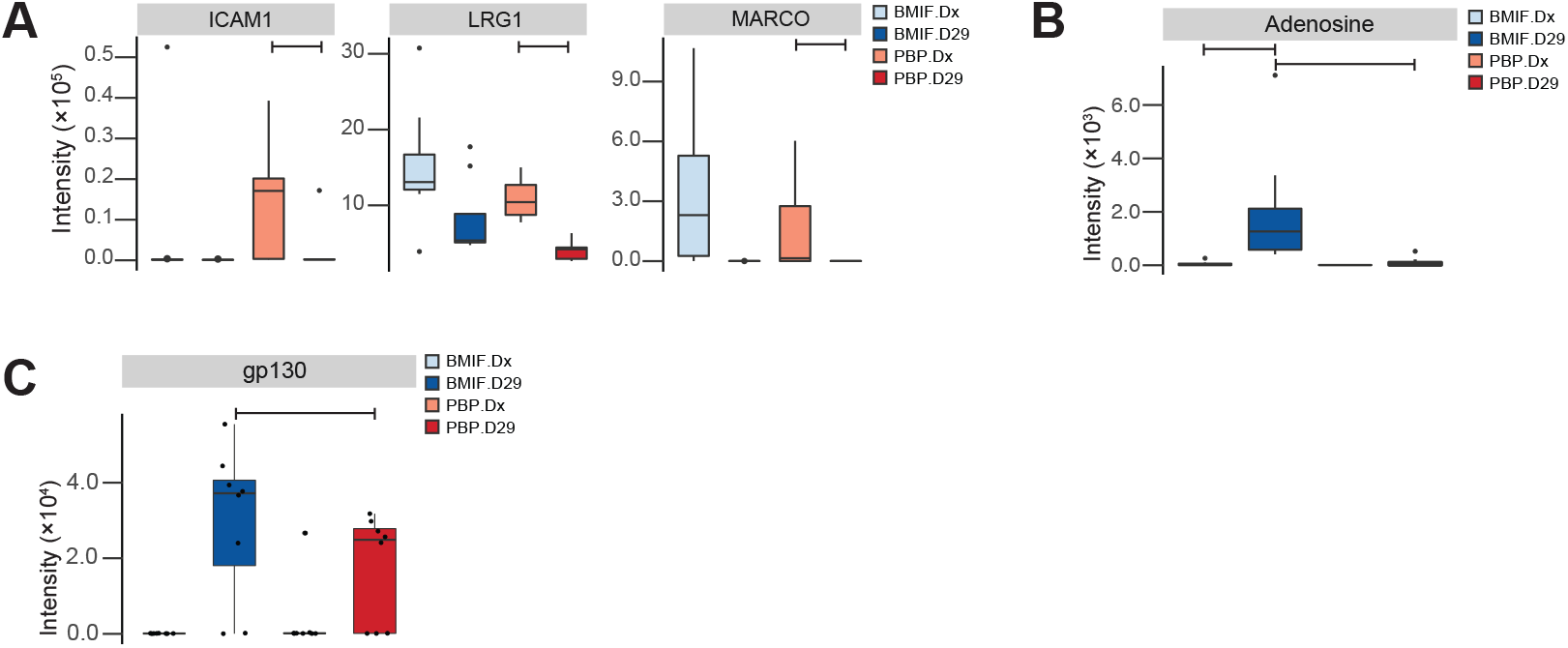
Immunosuppressive microenvironment in the after induction therapy. **(A)** Immune proteins with significant declines in intensity after induction therapy based on PBP differential analysis. **(B)** Intensity of adenosine across experimental groups. **(C)** Intensity of gp130/IL6ST across experimental groups. Bars indicate significant difference (limma, adjusted p-value ≤ 0.05) between experimental groups.

